# Another fly diuretic hormone: tachykinins increase fluid and ion transport by adult *Drosophila melanogaster* Malpighian ‘renal’ tubules

**DOI:** 10.1101/2024.02.22.581479

**Authors:** Marishia A. Agard, Meet Zandawala, Jean-Paul V. Paluzzi

## Abstract

Insects like the model organism *Drosophila melanogaster* must modulate their internal physiology to withstand changes in temperature and availability of water and food. Regulation of the excretory system by peptidergic hormones is one mechanism by which insects maintain their internal homeostasis. Tachykinins are a family of neuropeptides that have been shown to stimulate fluid secretion from the Malpighian ‘renal’ tubules (MTs) in some insect species, but it is unclear if that is the case in the fruit fly, *D. melanogaster*. A central objective of the current study was to examine the physiological role of tachykinin signaling in the MTs of adult *D. melanogaster*. Using the genetic toolbox available in this model organism along with *in vitro* and whole animal bioassays, our results indicate that *Drosophila* tachykinins (DTKs) function as diuretic hormones by binding to the DTK receptor (DTKR) expressed in stellate cells of the MTs. Specifically, DTK activates cation and anion transport across the stimulated MTs, which impairs their survival in response to desiccation due to their inability to conserve water. Thus, besides their previously described roles in neuromodulation of pathways controlling locomotion and food search, olfactory processing, aggression, lipid metabolism and metabolic stress, processing of noxious stimuli and hormone release, DTKs also appear to function as *bona fide* endocrine factors regulating the excretory system and appear essential for the maintenance of hydromineral balance.

**Summary statement:** *Drosophila* tachykinins are diuretic hormones in the fly that regulate activity of the MTs and, consequently, contribute towards the maintenance of ionic and osmotic homeostasis.

## Introduction

Insects like the fruit fly, *Drosophila melanogaster*, are routinely faced with challenges in their natural environment including changes to temperature, humidity, food availability and access to water, which necessitate feedback mechanisms and other physiological processes to maintain homeostasis (Chown et al., 2011; Leopold and Perrimon, 2007). Insects living in arid habitats have evolved to increase their tolerance to desiccation by reducing the rate of water loss and conserving body water content (Chown et al., 2011). One strategy that insects utilize to enhance survival in response to environmental stresses involves the regulation of the excretory organs, composed of the Malpighian tubules (MTs) and hindgut (Benoit, 2010; Beyenbach et al., 2010). The MTs in *D. melanogaster* are composed of two main epithelial cell types: (1) the principal cells, which are the most abundant cells making up the MTs that mediate transepithelial secretion of Na^+^ and K^+^ ions using the ionomotive enzyme V-type H^+^-ATPase; and (2) the relatively less abundant stellate cells that are responsible for Cl^−^ and water transport (Beyenbach et al., 2010; Cabrero et al., 2020). Hormones that act upon the MTs can be released from neurosecretory cells or endocrine cells localized in the central nervous system or peripheral tissues (e.g. intestines), respectively (Hartenstein, 2006). These hormones target distinct epithelial cell types of the MTs to function as either diuretic or anti-diuretic factors, mainly by interacting with specific G protein-coupled receptors (GPCRs) that can stimulate or inhibit transepithelial ion transport and, consequently, osmotically obliged water (Audsley et al., 1992; Hartenstein, 2006; Terhzaz et al., 1999). Therefore, to overcome excessive or limited amounts of water and solutes associated with their particular diet or environment, *Drosophila* modulate their MTs and other components of the excretory system by peptidergic hormones and other neurochemicals to regulate primary urine production and re-absorptive processes (Coast et al., 2002).

Tachykinins are a group of neuropeptides that are expressed in the central nervous system and intestine in both vertebrates and invertebrates and are widely considered as pleiotropic factors (Nässel et al., 2019). Insect tachykinin-related peptides have been shown to function as myostimulatory peptides on peripheral tissues including the foregut, midgut, hindgut, and oviduct (Schoofs et al., 1990a; Schoofs et al., 1990b). Studies have shown that insect tachykinins, such as those found in locusts and cockroaches, display myotropic effects as determined via muscle contraction assays (Muren and Nässel, 1996; Winther et al., 1998). One prototypical insect tachykinin-related peptide, locustatachykinin, has also been shown to stimulate tubule writhing activity, which may assist with urine flow (Coast, 1998). In addition to their myotropic effects, exogenous tachykinins stimulate secretion by the MTs in the tobacco hornworm, *Manduca sexta* (Skaer et al., 2002). Similarly, locustatachykinin has been shown to elicit dose-dependent diuretic effects in isolated tubules from two locust species, *Schistocerca gregaria* and *Locusta migratoria* (Johard et al., 2003). These effects of tachykinins and tachykinin-related peptides can be mediated via both cAMP and PLC-IP_3_-calcium signal transduction pathways (Birse et al., 2006; Torfs et al., 2000).

We ask here whether *Drosophila* tachykinins (DTKs), act as diuretic (neuro)peptide hormones with a potential role in regulating excretory organs that contribute towards ion-water balance and homeostasis. The *Drosophila tachykinin* (*Dtk)* gene is expressed in interneurons of the central nervous system including in the brain and subesophageal ganglion (Siviter et al., 2000). DTKs are localized in about 150 neurons in the adult fly central nervous system aiding in olfactory and locomotor behaviour in search of food during times of hunger (Winther et al., 2003; Winther et al., 2006). DTKs are also co-expressed with ion-transport peptide (ITP), and short neuropeptide F (sNPF) in neurosecretory cells in the adult *Drosophila* brain, where they may function as neuromodulators (Kahsai et al., 2010). Studies have also revealed that DTKs are expressed in enteroendocrine cells in the midgut where they regulate physiological functions including intestinal lipid metabolism (Siviter et al., 2000; Song et al., 2014). *Dtk* encodes for six structurally related peptide isoforms (DTK-1-6) that bind and activate two GPCRs, *Drosophila* tachykinin receptor (DTKR or TkR99D) and neurokinin K receptor (NKD or TkR86C) (Li et al., 1991; Monnier et al., 1992; Siviter et al., 2000). While NKD has been detected in the central nervous system and midgut (Jiang et al., 2013; Poels et al., 2009), some evidence indicated that NKD is not a tachykinin receptor. Instead, it has more recently been recognized as a *bona fide* receptor of natalisin, which is a tachykinin-related neuropeptide family in arthropods that influences reproductive physiology in insects (Jiang et al., 2013). Comparatively, DTKR is expressed in the central nervous system, midgut, and MTs and is activated by DTK-1-5 (Birse et al., 2006). However, recent studies have revealed that DTKs from a certain subset of brain neurons target neurons that express both receptors, DTKR and NKD, to function as neuromodulators of aggressive behaviour and visual aversive responses (Tsuji et al., 2023; Wohl et al., 2023). In the current study, we focused on the potential function of DTKs in regulating transepithelial secretion by the adult *D. melanogaster* MTs via DTKR.

Here, we confirm the cell-specific expression of DTKR in the fly MTs and provide insight into the physiological role of DTK/DTKR signaling in relation to ion-water balance. Based on earlier reports examining tachykinin activity on insect MTs (Johard et al., 2003; Skaer et al., 2002; Vanden Broeck et al., 1999; Winther and Nässel, 2001), we predicted that fly DTKs function as diuretic hormones by activating DTKR expressed in the stellate cells of the MTs to stimulate fluid transport, which may be important for the maintenance of hydromineral balance, especially during times of stress. To address these hypotheses, we first conducted a single-nucleus RNA sequencing (snRNA-seq) analysis of *Drosophila* MTs to determine the expression of DTKR (TkR99D). We then used two independent cell-specific driver lines to knock down *DTKR* expression in the MTs and measured the transcript levels of *DTKR* in this organ of the adult fruit fly. To further establish the physiological role of DTK/DTKR signaling in the MTs, various *in vitro* assays were conducted on isolated MTs along with whole animal stress assays where DTKR was specifically knocked down in MTs. Our results strongly support the notion that DTKR is indeed expressed in the stellate cells, since knockdown in the MTs affects renal function in the adult fruit fly that ultimately disrupts ion transport capacity. Finally, knockdown of DTKR exclusively in stellate cells of the MTs leads to increased lifespan during desiccation stress, which corroborates the diuretic activity of DTKs on the MTs and their influence on hydromineral homeostasis. Thus, these findings indicate that DTK/DTKR signaling contributes towards ionic and osmotic balance as DTKR knockdown flies displayed lower ion secretion and enhanced survival during desiccation stress. Overall, these findings establish the role of DTKs in hormonal control of the renal organs and build upon a growing list of diuretic and anti-diuretic factors in *Drosophila* that are fundamental for hydromineral homeostasis.

## Materials and Methods

### Fly lines and husbandry

Fly lines were maintained at room temperature with a 12hr:12hr light:dark cycle. The wild-type (*w^1118^*) flies, GAL4 and UAS fly lines used in this study are listed in Table 1. All fly lines were reared in vials containing food prepared as follows: 100 g/L sucrose, 50 g/L yeast, 12 g/L agar, 3 ml/L propionic acid, and 1.5 g/L nipagin. All experiments utilized 5-7 day-old (post-eclosion) adult females.

**Table 1.** Fly lines used in the study.

| Fly lines | Inserted on chromosome | Role | Source or reference | Stock number |
| --- | --- | --- | --- | --- |
| $w^{1118}$ (Wild-type) | ----- | Wild-type (control) | BDSC | 5905 |
| TkR99D (DTKR) GAL4 | III | Expresses GAL4 in TkR99D pattern. | BDSC | 76208 |
| *uro*-GAL4 (PC GAL4) | II | Expresses GAL4 in the pattern of the urate oxidase gene in principal cells of the Malpighian tubules | Dr. Kenneth Halberg, (Halberg et al., 2016) | ----- |
| *c724*-GAL4 (SC GAL4) | II | Expresses GAL4 in the stellate cells of the Malpighian tubules | Dr. Laura Nilson, (Feingold et al., 2019) | ----- |
| UAS-*DTKR*-RNAi (RNAi 1) | III | UAS-*DTKR*-RNAi 1<sup>st</sup> line that encodes for the *DTKR*-RNAi construct | BDSC | 27513 |
| UAS-*DTKR*-RNAi (RNAi 2) | II | UAS-*DTKR*-RNAi 2<sup>nd</sup> line that encodes for the *DTKR*-RNAi construct | BDSC | 55732 |
| JFRC81-10xUAS-IVS-Syn21-GFP-p10 (GFP) | ----- | GFP is strongly expressed in cells where GAL4 has been expressed. | (Zandawala et al., 2021) | ----- |

### Single-nucleus RNA-sequencing Analysis

Single-nucleus transcriptomes of Malpighian tubules were mined using the FlyCellAtlas dataset generated earlier (Xu et al., 2022). The parameters used to identify Tkr99D positive stellate cells were: TkR99D > 0 & annotation == “adult Malpighian tubule stellate cell of main segment”. All analyses were performed in R-Studio (v2022.02.0) using the Seurat package (v4.1.1 (Hao et al., 2021)). Heatmaps were constructed using the pheatmap package. Code available upon request.

### Fluorescence microscopy

Fluorescence microscopy was performed to visualize GFP expression in the MTs. The JFRC81-10xUAS-IVS-Syn21-GFP-p10 (GFP) fly line was crossed with the following GAL4 lines: *TkR99D* (*DTKR*)-GAL4, *uro*-GAL4, and *c724*-GAL4. The MTs were dissected and isolated from the transgenic flies in phosphate-buffered saline (PBS). The guts were then fixed with 4% paraformaldehyde for 1 hour on the rocker at room temperature followed by three washes each for 10-15 minutes with PBS. The guts were mounted in 50% glycerol and visualized with the EVOS FL Auto live-cell imaging system (Life Technologies, Burlington, ON).

### RT-qPCR

RT-qPCR was used to measure knockdown efficiency of DTKR by RNAi in fly MTs. Total RNA was extracted from 30-35 sets of MTs isolated from 5-7-day old female flies in five independent biological replicates using the Monarch Total RNA Miniprep kit (New England BioLabs, Whitby, ON, Canada). Total RNA was quantified using a UV spectrophotometer (Synergy 2 microplate reader, BioTek, Winooski, VT) and cDNA synthesis was performed using the iScript Reverse Transcription Supermix (Bio-Rad, Mississauga, ON, Canada). RT-qPCR was set up with quadruplicate technical replicates using PowerUP™ SYBR® Green Master Mix (Applied Biosystems, Carlsbad, CA, USA). The template cDNA was diluted 5-fold and used for RT-qPCR using a StepOnePlus Real-Time PCR System (Applied Biosystems, Carlsbad, CA, USA) that ran for 40 cycles: denaturation (95(), annealing (60(), and extension (72(). Forward (GAGTAAGCGAAGGGTGGTGAAG) and reverse (GAACGGCAGCCAGCAGAT) gene-specific primers described previously (Birse et al., 2011; Ignell et al., 2009) were used to quantify DTKR expression in the MTs. Ribosomal protein L32 (RpL32) and alpha-tubulin were used as the reference genes for the RT-qPCR, as previously described (Ponton et al., 2011). Standard curves were obtained using serial dilutions of the MTs cDNA template from female *w^1118^* yielding primer efficiencies ranging from 90-103%. For each replicate, the ΔΔCt method was used to determine the relative quantification (RQ) value, which was normalized to the geometric mean of RpL32 and alpha-tubulin reference genes (Sajadi et al., 2020). Statistical analysis was performed on the log-transformed data.

### Malpighian tubules secretion assay

The fluid secretion assay was adapted from Ramsay (Ramsay, 1954). Female adult fruit flies were isolated and dissected in *Drosophila* saline (mmol l^−1^): 10 glutamine, 20 glucose, 15 MOPS, 4.3 NaH_2_PO_4_, 10.2 NaHCO_3_, 8.5 MgCl_2_ (hexahydrate), 2 CaCl_2_ (dihydrate), 20 KCl, 117.5 NaCl that was titrated to pH 7 (Vanderveken and O’Donnell, 2014). The MTs were placed into small wells in a Sylgard-lined Petri dish containing 20µL of the DTK treatment in a 1:1 ratio of *Drosophila* saline: Schneider’s Insect Medium (Sigma-Aldrich, Oakville, ON). The MTs were treated with 10^−4^-10^−13^ M DTK-1 (APTSSFIGMR-amide) to determine the full dose-response curve and 10^−7^ M DTK-1 (∼EC_50_) was the selected dose to test out secretion rates from knockdown flies as identical secretion rates were observed when MTs were treated with the same concentration of other DTK isoforms, including DTK-2 and DTK-3. The proximal end of the tubule was wrapped around a minutien pin to measure the urine droplet from the MTs after a 60 min incubation. After the treatment period, the urine droplet was measured using a calibrated eyepiece micrometer from the dissecting microscope to calculate the fluid secretion rate (nL min^−1^) with the following equation: secretion 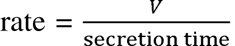, where secretion time refers to the secretion time 60-minute incubation, and *V* refers to the volume of the secreted droplet.

### Use of Ion-selective microelectrodes (ISME) to measure ion concentration from isolated tubules

Ion-selective microelectrodes (ISME) were used to measure Na^+^, K^+^, and Cl^−^ concentrations from MTs isolated from wild-type (*w^1118^*) and transgenic flies treated with DTK-1 (treatment) and 0.5% DMSO (control as DTK stocks were made up in DMSO). Glass capillaries for microelectrodes (TW150-4, World Precision Instruments, Sarasota, FL, USA) and reference electrodes (1B200F-4, World Precision Instruments, Sarasota, FL, USA) were prepared using a Sutter P-97 Flaming Brown pipette puller (Sutter Instruments, San Raffael, CA, USA). The ion-selective microelectrodes were silanized using N, N-dimethyltrimethylsilylamine (Sigma-Aldrich, Oakville, ON) as previously described (Lajevardi et al., 2021). The reference electrode was backfilled with 500 mmol l^−1^ KCl. The Na^+^-selective microelectrode was backfilled with 100 mmol l^−1^ NaCl, while the K^+^-selective microelectrode was backfilled with 100 mmol l^−1^ KCl, and the Cl^−^-selective microelectrode was backfilled with 150 mmol l^−1^ KCl. The tip of the ion-selective microelectrodes were forward-filled with an ion-specific ionophore: Na^+^ ionophore (sodium ionophore II cocktail A, Fluka), K^+^ ionophore (potassium ionophore I cocktail B, Fluka), and Cl^−^ ionophore (Chloride ionophore I - cocktail A, Sigma-Aldrich). The tips of the electrodes were coated with polyvinyl chloride dissolved in tetrahydrofuran (Sigma-Aldrich) to prevent the movement of the ionophore when submerged in the mineral oil as described previously (Lajevardi et al., 2021).

The silver chloride wires from the electrometer were inserted into the microelectrodes so that the differential voltage could be measured and recorded with the electrometer and a data acquisition system (Picolog for Windows, version 5.25.3). The ISME was calibrated in the following standard solutions: Na^+^ microelectrode, 200 mmol l^−1^ NaCl and 20 mmol l^−1^ NaCl +180 mmol l^−1^ LiCl; K^+^ microelectrode, 150 mmol l^−1^ KCl and 15 mmol l^−1^ KCl + 135 mmol l^−1^ LiCl; Cl^−^ microelectrode, 150 mmol l^−1^ KCl and 15 mmol l^−1^ KCl. The ISME and reference electrodes were positioned into the secreted droplet using the micromanipulators. The ion concentration was calculated using the equation previously described (Donini et al., 2008; Paluzzi et al., 2012; Rheault and O’Donnell, 2004): 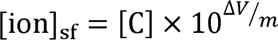, where [C] is the concentration of the calibration solution in mmol l^−1^, ΔV is the difference between the voltage recorded in the secreted fluid droplet and the voltage recorded in the calibration solution, and *m* is the voltage difference between the two standard calibrations (slope). Since K^+^ is known to interfere with the response of the Na^+^ microelectrode (based on the sodium ionophore II cocktail A), measurements of Na^+^ were corrected using the Nicolsky Eisenman equation and a selectivity coefficient of the Na^+^ microelectrode for K^+^ (K_NaK_ = 0.398) as previously described (Ammann and Anker, 1985; Ianowski and O’Donnell, 2004). Finally, the ion transport rates were calculated by multiplying the fluid secretion rates by the [ion]_sf_.

### Stress resistance assays

The stress assays were performed on 5-7 day-old female flies. For each of the four biological replicates, 15-20 flies were selected, and the number of surviving flies was recorded every 8 hours until all flies were dead. The desiccation assay consisted of placing 5-7 day-old female DTKR-RNAi flies and control flies into empty vials to deprive flies of food and water to induce desiccation (Liao et al., 2020; Söderberg et al., 2011). The starvation assay consisted of placing 5-7 day-old female DTKR-RNAi flies and control flies into vials containing 1% aqueous agar (Fisher Scientific, USA) (Liao et al., 2020; Söderberg et al., 2011).

### Statistical Analyses

All experimental data was graphed as mean±SEM. The data were analyzed using GraphPad Prism Version 8.0.0 (San Diego, California USA) to complete the appropriate statistical test such as the unpaired t-test, one-way ANOVA & two-way ANOVA with the addition of the post-hoc test (Tukeys and Šídák’s). The Log-rank (Mantel-Cox) test was used to determine significant differences in survival for stress assays after correcting for multiple comparisons by computing the Bonferroni-corrected significance level (a_Bonferroni_ = 0.025), which considers the family-wise alpha level (p = 0.05) and the number of comparisons.

## Results

### Single-nucleus RNA sequencing analysis of MTs confirms DTKR expression in the stellate cells

Gene expression analysis was conducted on an existing snRNA-seq dataset to confirm the cell-type specific expression of DTKR (TkR99D) in the MTs (Xu et al., 2022). The MTs are composed of 10 distinct clusters, which can express a variety of diuretic (or anti-diuretic) hormone receptors. Based on the snRNA-seq analysis, a number of diuretic hormone receptors (Cannell et al., 2016; Coast et al., 2002; Johnson et al., 2004; Radford et al., 2002a) including *TkR99D*, *leucokinin receptor* (*Lkr*), and *diuretic hormone 44 receptor 2* (*DH44-R2*) are highly expressed in the cluster comprised of the stellate cells of the main segments (**Fig. 1A-C**). Specifically, roughly 75% of the stellate cells of the main segment express *TkR99D*, *Lkr* and *DH44-R2*. In contrast, distinct peptidergic regulators of the MTs have receptors that are not expressed in stellate cells including those for diuretic hormone 31 (DH_31_) and Capa peptides, but instead display high expression in clusters representing the principal cells (**Fig. 1A-C**) in strong agreement with earlier findings (Johnson et al., 2005; Terhzaz et al., 2012). Notably, the *tyramine receptor* (*TyrR*) was strongly expressed in both the principal cells of the lower segment and stellate cells of the main segments (**Fig. 1A-C**). Intriguingly, *TkR99D* is only expressed in the star-shaped (main segment) stellate cells while the *Lkr* is expressed in both bar (initial segment) and star-shaped (main segment) stellate cells (**Fig. 1C**). Interestingly, similarity clustering of stellate cells based on expression of diuretic hormone receptors shows that cells expressing *Tkr99D* also express *Lkr* (**Fig. 1D**). Independently, TkR99D-GAL4 drives strong GFP expression in the star-shaped stellate cells, thus validating our snRNA-seq analysis (**Fig. 1E**). Further examination of *TkR99D*-positive stellate cells shows that they express a variety of diuretic hormone receptors and water channels (*Drip and Prip*), the latter of which might be central for primary urine production when tubules are stimulated by DTKs (**Fig. 1F**).

**Fig. 1.**
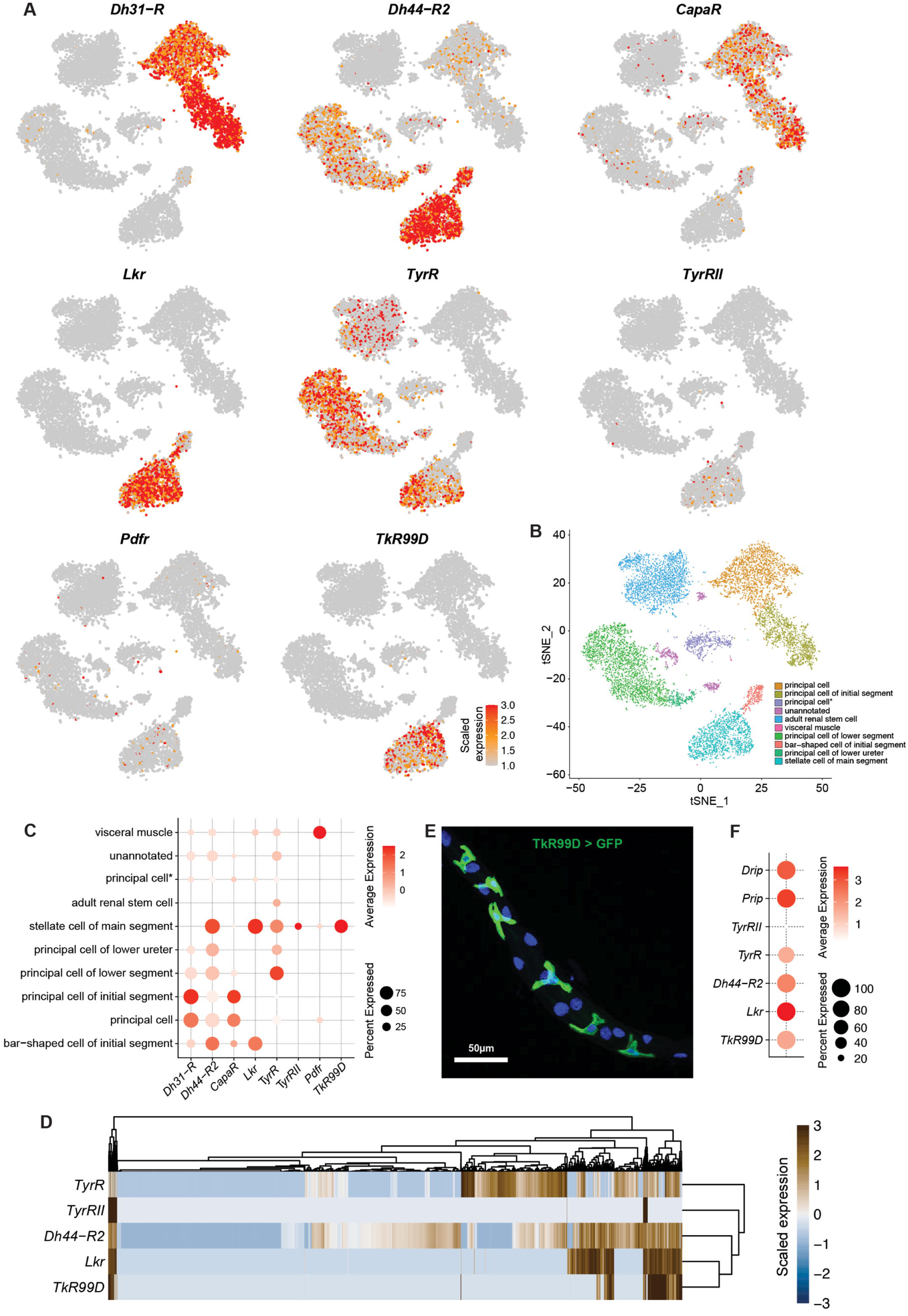
Single-nucleus RNA sequencing analysis distinguishing the Malpighian tubules cell types and expression of hormone receptors and channels. (A) Expression of hormone receptors in single cell transcriptomes of Malpighian tubules. (B) Clustering of different cell types in Malpighian tubules. (C) Dotplot showing expression of hormone receptors in different cell types of Malpighian tubules. (D) Heatmap of all stellate cells showing expression of hormone receptors. Leucokinin receptor (Lkr) and Tkr99D show overlapping expression pattern. (E) Tubules isolated from TkR99D>JFRC81-10xUAS-IVS-Syn21-GFP-p10 (GFP) flies displayed high GFP expression in the star-shaped stellate cells (scale bar: 50µm). (F) Dot plot showing expression of diuretic hormone receptors and aquaporins in Tkr99D positive cells.

### Examining the physiological role of DTK on isolated MTs

Secretion rates were measured from isolated MTs using the classic *in vitro* assay described by Ramsay to determine the full dose-response characteristics of DTK-1, which was used as a representative of all *Dtk* derived peptides since all peptide isoforms have similar efficacy in DTKR activation (Birse et al., 2006; Ramsay, 1954). MTs were isolated from wild-type (*w^1118^*) flies and treated with different concentrations of DTK-1 (ranging from 10^−4^ to 10^−13^ M) to determine an intermediate concentration that could then be used on MTs isolated from DTKR-RNAi flies since high doses might not be as sensitive to reduced receptor expression. The dose-response analysis demonstrates the diuretic activity of DTK-1, with an EC_50_ of 1.126 x 10^−7^ M (**Fig. 2A**), which is similar to the activity of other insect tachykinins such as locustatachykinin I tested on *Locusta migratoria* and *Schistocerca gregaria* MTs (Johard et al., 2003). Indeed 10^−7^ M DTK-1 was shown to be an intermediate dose as it elicited significantly higher fluid secretion rates compared to unstimulated and 10^−9^ M DTK-1 treated MTs while displaying significantly lower rates of secretion when compared to 10^−5^ M DTK-1 treated tubules (**Fig. 2B**). In addition, the MTs were treated separately with DTK-1, DTK-2, and DTK-3 at 10^−7^ M, which revealed similar secretion rates and diuretic activity (**Fig. 2C**), indicating that multiple DTK peptide isoforms have similar effects in driving secretion by isolated tubules.

**Fig. 2.**
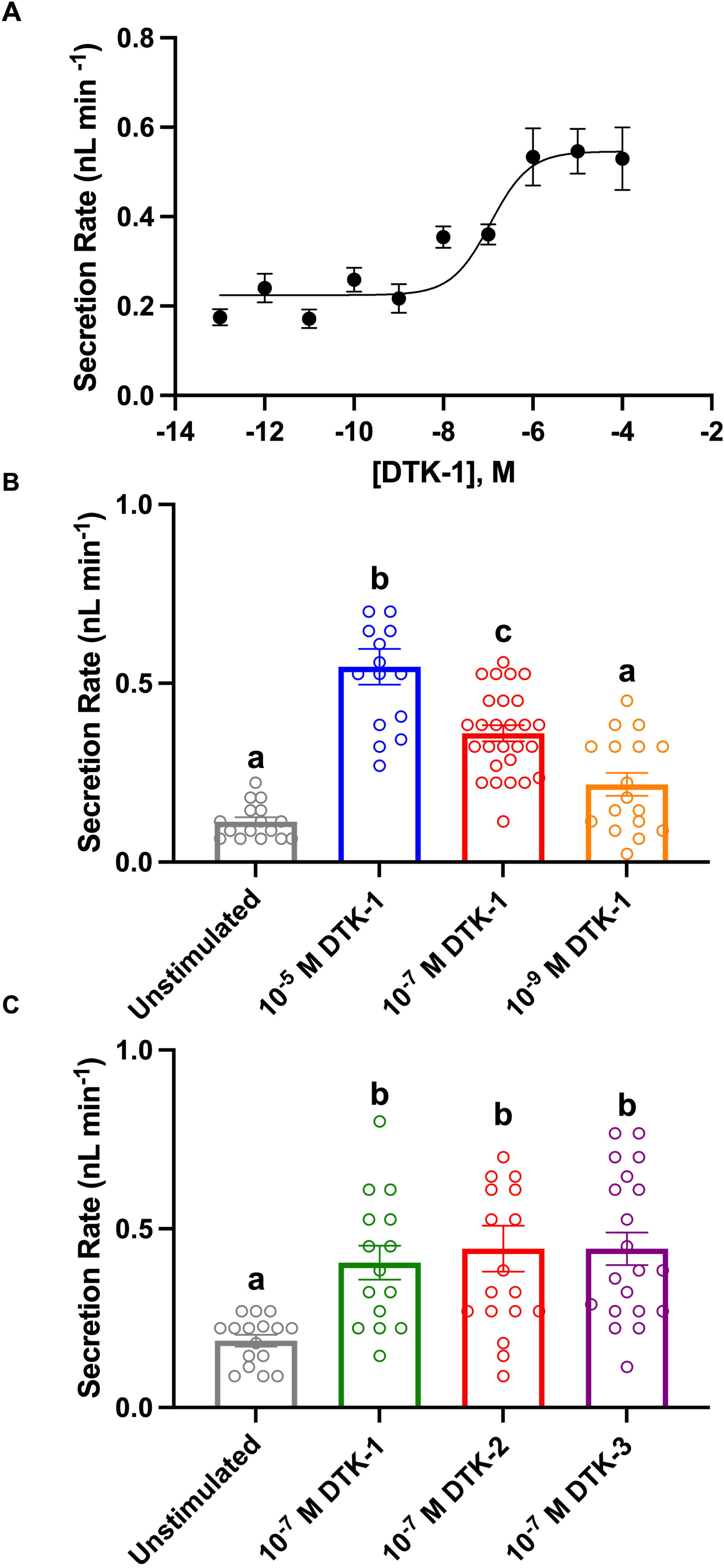
Secretion rates by *in vitro* isolated Malpighian tubules in response to different concentrations of *Drosophila* tachykinin-1 (DTK-1) The MTs isolated from wild-type (WT) flies (A) displayed a dose-dependent activation of fluid secretion as shown by the activity of various concentrations of DTK-1, where the EC_50_ was 1.126 x 10^−7^ M DTK-1 (assessed by log(agonist) vs. response (three parameters) non-linear regression). (B) Tubules treated with 10^−7^ M DTK-1 displayed significant stimulation of secretion compared to unstimulated and 10^−9^ M DTK-1 treatment, while displaying significantly lower secretion rates compared to 10^−5^ M DTK-1 (p<0.05, assessed by One-way ANOVA). (C) Tubules displayed identical significant increases in fluid secretion rates when treated with DTK-1, DTK-2, and DTK-3 at 10^−7^ M compared to unstimulated MTs (p<0.05, assessed by One-way ANOVA).

### Knockdown of DTKR expression in adult fly MTs

To functionally validate the expression of DTKR in the adult fruit fly MTs, we utilized the bipartite GAL4-UAS system to drive *DTKR* RNAi using two independent cell-specific GAL4 drivers: (1) *uro*-GAL4 and (2) *c724*-GAL4, which drive GFP expression in principal and stellate cells, respectively (**Fig. 3A,B**). Previous studies have established the expression patterns of these two lines with *uro*-GAL4 driving expression in the more abundant principal cells found along the length of both the anterior and posterior MTs, including the initial and the main segments (Sözen et al., 1997; Weaver et al., 2020). Comparatively, *c724*-GAL4 drives expression in the star-shaped stellate cells from the initial to the main segment of both anterior and posterior MTs (Sözen et al., 1997). First, the principal cell *uro*-GAL4 driver line was crossed with two different RNAi lines (*uro*-GAL4>UAS-*DTKR* RNAi 1 and *uro*-GAL4>UAS *DTKR* RNAi 2) in an attempt to knock down *DTKR* in the principal cells of the MTs. Tubules from the experimental flies displayed a similar *DTKR* abundance to the controls (**Fig. 3C**) with no evidence of a reduction in *DTKR* transcript levels. Next, the stellate cell driver was crossed with the two DTKR-RNAi lines (*c724*-GAL4>UAS-*DTKR* RNAi 1 & *c724*-GAL4>UAS-*DTKR* RNAi 2) to knock down *DTKR* in stellate cells of the MTs (**Fig. 3D**). In contrast to the principal cell manipulation, a significant reduction in *DTKR* abundance was observed in MTs isolated from stellate cell driven DTKR-RNAi flies relative to MTs from controls that displayed normal DTKR expression (**Fig. 3D**). These results support the notion that DTKR is expressed in the stellate cells, but not the principal cells of the MTs, and hence identify the former cell type in MTs as the cellular target of circulating DTK. These results are consistent with the snRNA-seq data from the adult fruit fly that detected high expression of *DTKR* in the main segment stellate cells (**Fig. 1A,C,E**) of the MTs (Xu et al., 2022). This evidence also indicates that *DTKR* is co-expressed in stellate cells with other diuretic hormone receptors (**Fig. 1F**), including *Lkr* and *TyrR* (Blumenthal, 2003; Cabrero et al., 2013; Radford et al., 2002b).

**Fig. 3.**
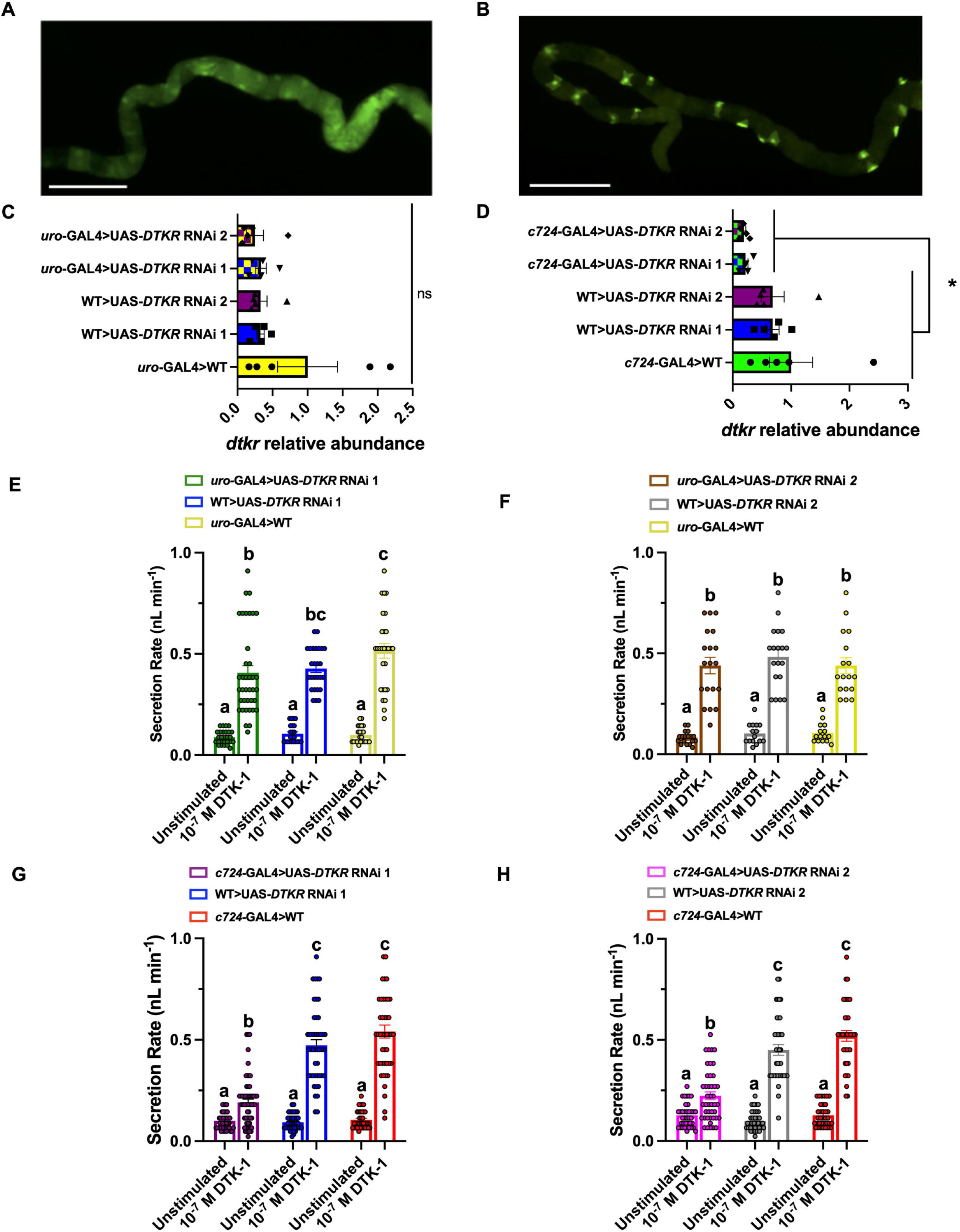
*DTKR* transcript levels in Malpighian tubules and fluid secretion rates isolated from different crosses to assess *DTKR* knockdown using either a principal or stellate cell GAL4 driver. (A) Tubules isolated from *uro*-GAL4>JFRC81-10xUAS-IVS-Syn21-GFP-p10 (GFP) flies displayed high GFP expression in the principal cells (scale bar: 400µm). (B) High GFP expression in the stellate cells of the tubules isolated from *c724*-GAL4>GFP flies (scale bar: 400µm). (C) Malpighian tubules isolated from flies where knockdown was driven by the principal cell *uro*-GAL4 displayed no changes in *DTKR* transcript levels. (D) Tubules isolated from flies where knockdown was driven by the stellate cell *c724*-GAL4 displayed a significant reduction in *DTKR* abundance. Similar *DTKR* knockdown in stellate cells was achieved using two different *DTKR* RNAi lines (p<0.05, assessed by two-way ANOVA). (E-F) Tubules isolated from knockdown flies driven by the *uro*-GAL4 displayed a significant increase in fluid secretion rates in response to DTK-1 compared to tubules from control crosses (p<0.05, assessed by two-way ANOVA). (G-H) Tubules isolated from *c724*-GAL4 and two UAS-*DTKR* RNAi crosses displayed significantly reduced stimulated secretion rates in response to DTK-1 treatment compared to tubules from control crosses (p<0.05, assessed by two-way ANOVA).

### Secretion rates from tubules isolated from knockdown flies

To examine the functional impact of *DTKR* knockdown in the renal tubules, the MT fluid secretion assay was completed on DTKR-RNAi flies in response to DTK-1 using the *in vitro* assay (Ramsay, 1954). Fluid secretion rates by MTs isolated from flies with DTKR-RNAi driven by the principal cell driver (*uro*-GAL4>UAS-*DTKR* RNAi 1 & *uro*-GAL4>UAS *DTKR* RNAi 2) and the genetic control flies displayed a significant increase (∼4-fold) in fluid secretion rates when treated with 10^−7^ M DTK-1 (**Fig. 3E,F**). The MTs from *uro*-GAL4 driven RNAi lines displayed similar secretion rates to the parental control lines under both basal (unstimulated) and DTK-1 stimulated conditions. These bioassay results support the RT-qPCR analysis and snRNA-seq data that indicate DTKR is not expressed in the principal cells of the MTs.

Fluid secretion rates were then measured in MTs isolated from flies with DTKR-RNAi driven by the stellate cell driver (*c724*-GAL4>UAS-*DTKR* RNAi 1 & *c724*-GAL4>UAS-*DTKR* RNAi 2) and the genetic control flies (**Fig. 3G,H**). Basal unstimulated fluid secretion rates by MTs from *c724*-GAL4>UAS-*DTKR* RNAi 1 & *c724*-GAL4>UAS-*DTKR* RNAi 2 flies were similar to parental control lines. However, the degree of DTK-1 stimulation relative to unstimulated MTs was 81-84% lower in *DTKR* knockdown flies compared to MTs from parental control lines. Thus, the knockdown of DTKR in stellate cells drastically decreases stimulated secretion rates in response to DTK-1, but rates were marginally (and significantly) elevated compared to the unstimulated basal rates of secretion. Altogether, the results confirm that reduced expression of DTKR specifically in stellate cells leads to lower fluid secretion rates in response to DTK-1, which complements the earlier data demonstrating strong GFP expression in the star-shaped stellate cells driven by TkR99D-GAL4 (**Fig 1E**). Similarly, this functional data for DTK-1 on stellate cells substantiates the cell-type specific expression results observed for *DTKR* in the snRNA-seq (**Fig. 1A,C,F**) and RT-qPCR analysis (**Fig. 3C,D**).

### DTK-1 alters transepithelial ion transport by adult Malpighian tubules

Given that DTK elicits stimulatory effects on fluid secretion rates of MTs isolated *ex vivo* from wild-type flies, we predicted that ion concentrations (or their transport rate) may be altered in the secreted fluid since passive water movement is coupled to ion transport. Before investigating the effects on transepithelial transport across MTs isolated from DTKR knockdown flies, the activity of DTK-1 was examined in MTs from wild-type flies. Using ion-selective microelectrodes (ISME) to measure select ion concentrations in the primary urine droplets from MTs (Lajevardi et al., 2021), we first quantified Na^+^ concentration in the secreted fluid and its transport rate following treatment with DTK-1, which revealed no significant changes in Na^+^ concentration in the secreted fluid (**Fig. 4A**) but a significant increase in Na^+^ transport rates (**Fig. 4B**) compared to unstimulated MTs. There was a significant decrease in the K^+^ concentration (**Fig. 4C**) in the secreted fluid and a significant increase in K^+^ transport rate (**Fig. 4D**) when wild-type MTs were treated with DTK-1. Notably, wild-type MTs treated with DTK-1 demonstrated a significant increase in Cl^−^ concentration (**Fig. 4E**) in the secreted fluid and an increase in the transport rate of this anion (**Fig. 4F**) compared to unstimulated tubules. Therefore, the results indicate that DTK drives the movement of cation and anions, similar to the *Drosophila* leucokinin that displays effects on overall Na^+^, K^+^, Cl^−^ transport (Cabrero et al., 2014). Altogether, these data indicate that, along with increased fluid secretion rates, DTK stimulates an increase in cation and anion transport by targeting DTKR on stellate cells.

**Fig. 4.**
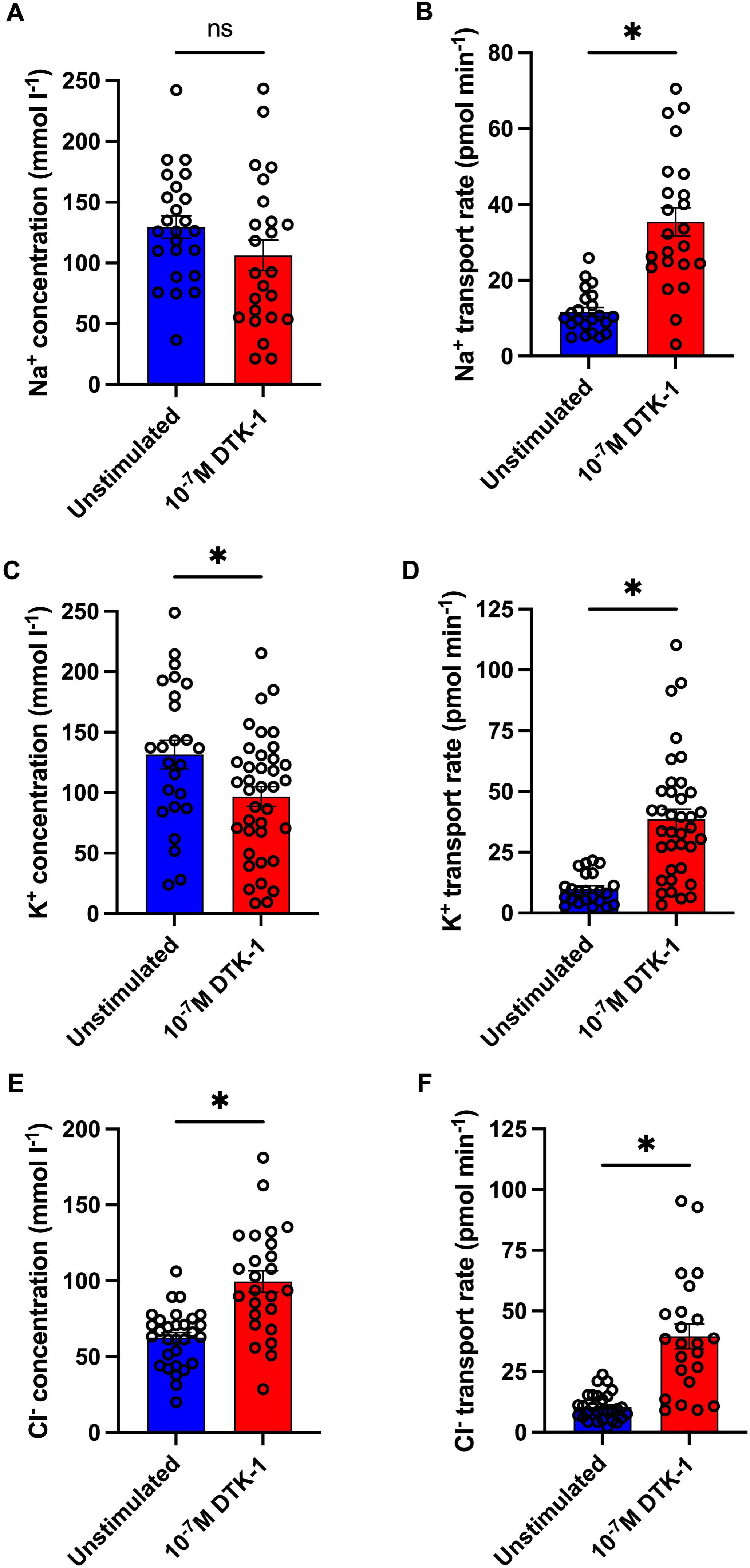
Ion concentration in the secreted fluid and transport rates from wild-type fly (WT) Malpighian tubules stimulated by *Drosophila* tachykinin-1. (A) Na^+^ concentration in the secreted fluid of tubules displayed no significant difference following DTK-1 treatment relative to unstimulated MTs (B) Significant increase in Na^+^ transport rate from tubules treated with DTK-1 (p<0.05, assessed by unpaired t-test). DTK-1 treatment resulted in significantly (C) lower K^+^ concentration in the secreted fluid and (D) elevated K^+^ transport rates by tubules (p<0.05, assessed by unpaired t-test). (E) Cl^−^ concentration in the secreted fluid and the (F) Cl^−^ transport rate significantly increased in MTs treated with DTK-1 (p<0.05, assessed by unpaired t-test).

We next examined how stellate cell-specific knockdown of DTKR in the MTs might impact transepithelial ion transport. In light of the significantly diminished DTK-1 stimulated fluid secretion in flies with *DTKR* knockdown in the stellate cells of the MTs using two different RNAi lines, we proceeded to use one of these (UAS-*DTKR* RNAi 1) for further analysis using ISME to examine the effect on tubule transepithelial ion transport. DTK-1 treatment did not affect Na^+^ concentration in the secreted fluid compared to unstimulated MTs, which was similarly observed in *DTKR* knockdown and control flies (**Fig. 5A**). Likewise, the K^+^ concentration in the secreted fluid did not change significantly following DTK-1 treatment compared to unstimulated MTs, and a similar trend was observed in knockdown and control flies (**Fig. 5B**). Notably, the Na^+^ and K^+^ transport rate by MTs in *DTKR* knockdown flies after treatment with DTK-1 was not different from unstimulated conditions while transport of both these cations significantly increased following DTK-1 treatment of MTs in control flies (**Fig. 5C,D**) consistent with data on wildtype flies (**Fig. 4B,D**). Comparatively, the Cl^−^ concentration in tubule secreted fluid and the transport rate of this anion was unchanged following DTK-1 treatment in *DTKR*-RNAi flies whereas significantly elevated Cl^−^ concentration and transport rate was observed following DTK-1 stimulation of MTs from control flies (**Fig. 5E,F**) in agreement with observations on wildtype flies (**Fig. 4E,F**). In summary, while there was an increase in the Cl-concentration in the secreted fluid and transport rate of this anion in control flies, *DTKR* knockdown abolished these activated epithelial transport characteristics.

**Fig. 5.**
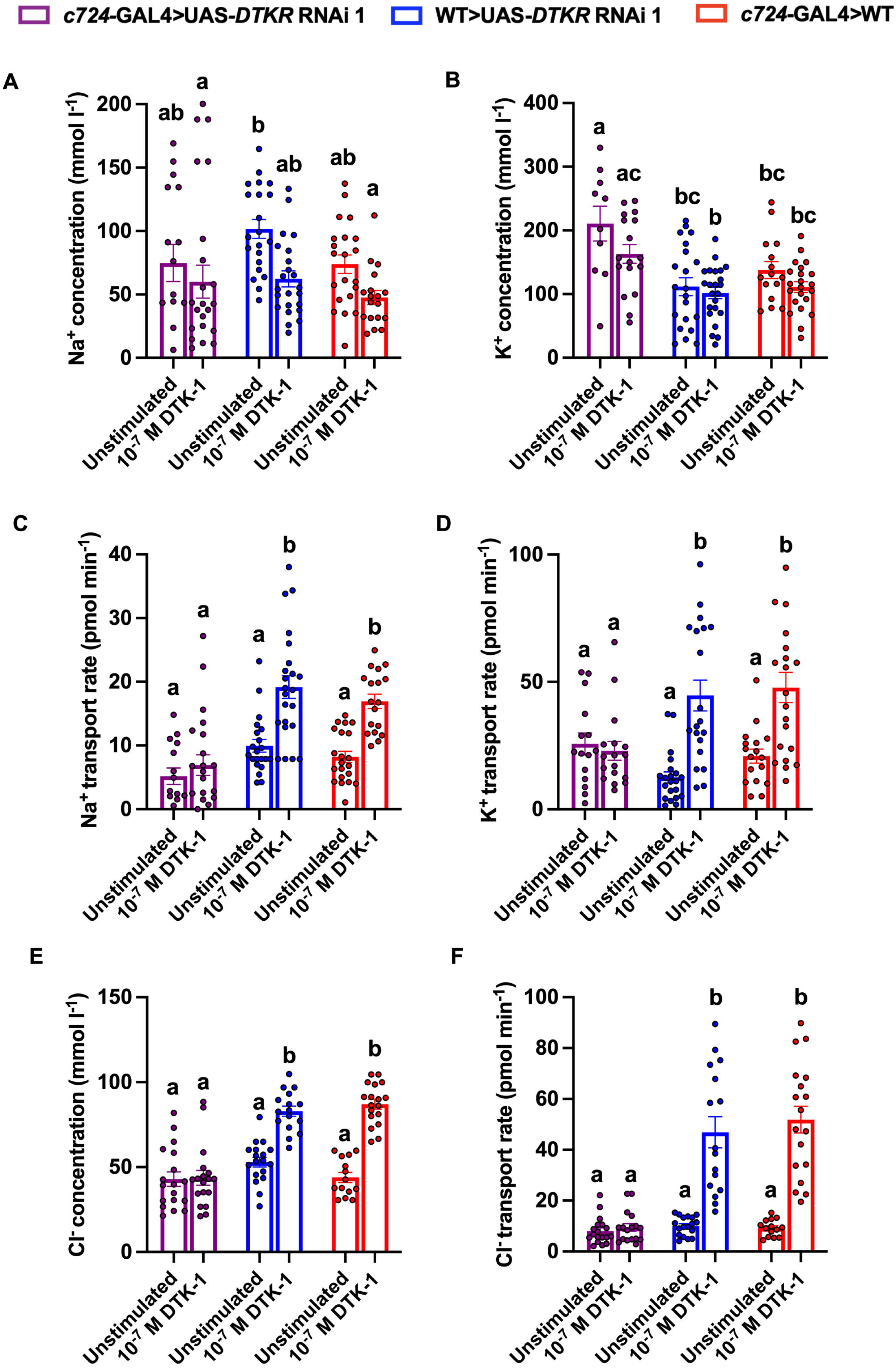
Effects of *Drosophila* tachykinin-1 on the ion concentration in secreted fluid and their transport rates following knockdown of *DTKR* in stellate cells. (A) Na^+^ and (B) K^+^ concentration in secreted fluid of the *DTKR* knockdown MTs was unchanged after DTK-1 treatment when compared to the unstimulated tubules and a similar trend was observed in MTs from control flies. (C) Na^+^ and (D) K^+^ transport rates were unchanged compared to unstimulated tubules following DTK-1 treatment in *DTKR* knockdown MTs, while transport rates of both cations increased significantly (> two-fold) following DTK-1 treatment in control flies. (E) Cl^−^ concentration in the secreted fluid was not significantly different compared to unstimulated MTs in *DTKR* knockdown following DTK-1 treatment while in Cl-concentration significantly increased in the secreted fluid of control fly MTs following DTK-1 treatment. Similarly, (F) Cl^−^ transport rate was unchanged compared to unstimulated tubules in *DTKR* knockdown MTs following DTK-1 treatment while transport of this anion increased significantly (> four-fold) in control fly MTs following DTK-1 treatment. All statistical comparisons were assessed by two-way ANOVA and Šídák’s post-hoc test with p<0.05 considered significant.

### Renal-specific DTKR knockdown improves desiccation and starvation survival

Considering our data revealing that *DTKR* knockdown in stellate cells reduces DTK-1 stimulated transepithelial transport, we predicted that these flies may also exhibit an increase in lifespan during stress conditions challenging fluid and ion homeostasis, such as desiccation. Stress assays were completed on DTKR-RNAi 1 flies driven by the stellate cell driver as it was the most effective in knocking down *DTKR* levels (**Fig. 1D**). Flies were deprived of water and food to induce desiccation stress, which revealed DTKR knockdown in stellate cells of the MTs (*c724*-GAL4>UAS-*DTKR* RNAi 1) resulted in a significant increase in lifespan with a maximal survival of 90 hours compared to either of the parental control crosses (WT>UAS-*DTKR* RNAi 1, p = 0.0047; *c724*-GAL4>WT, p < 0.0001) (**Fig. 6A**). Comparably, in the starvation stress assay where flies were only deprived of food, but not water (**Fig. 6B**), flies with DTKR knockdown in stellate cells survived significantly longer than WT>UAS-*DTKR* RNAi 1 controls (p<0.0001) while their survival did not differ compared to *c724*-GAL4>WT controls (p = 0.1255). Thus, based on the current data it is unclear whether DTK/DTKR renal signaling may be involved in feedback mechanisms related to nutritional stress and this should be further explored in future studies. Altogether, the results indicate that *DTKR* knockdown in the MTs may contribute towards greater water conservation in the animal due to decreased diuretic activity, which in turn enhances their dehydration tolerance and survival during desiccation stress.

**Fig. 6.**
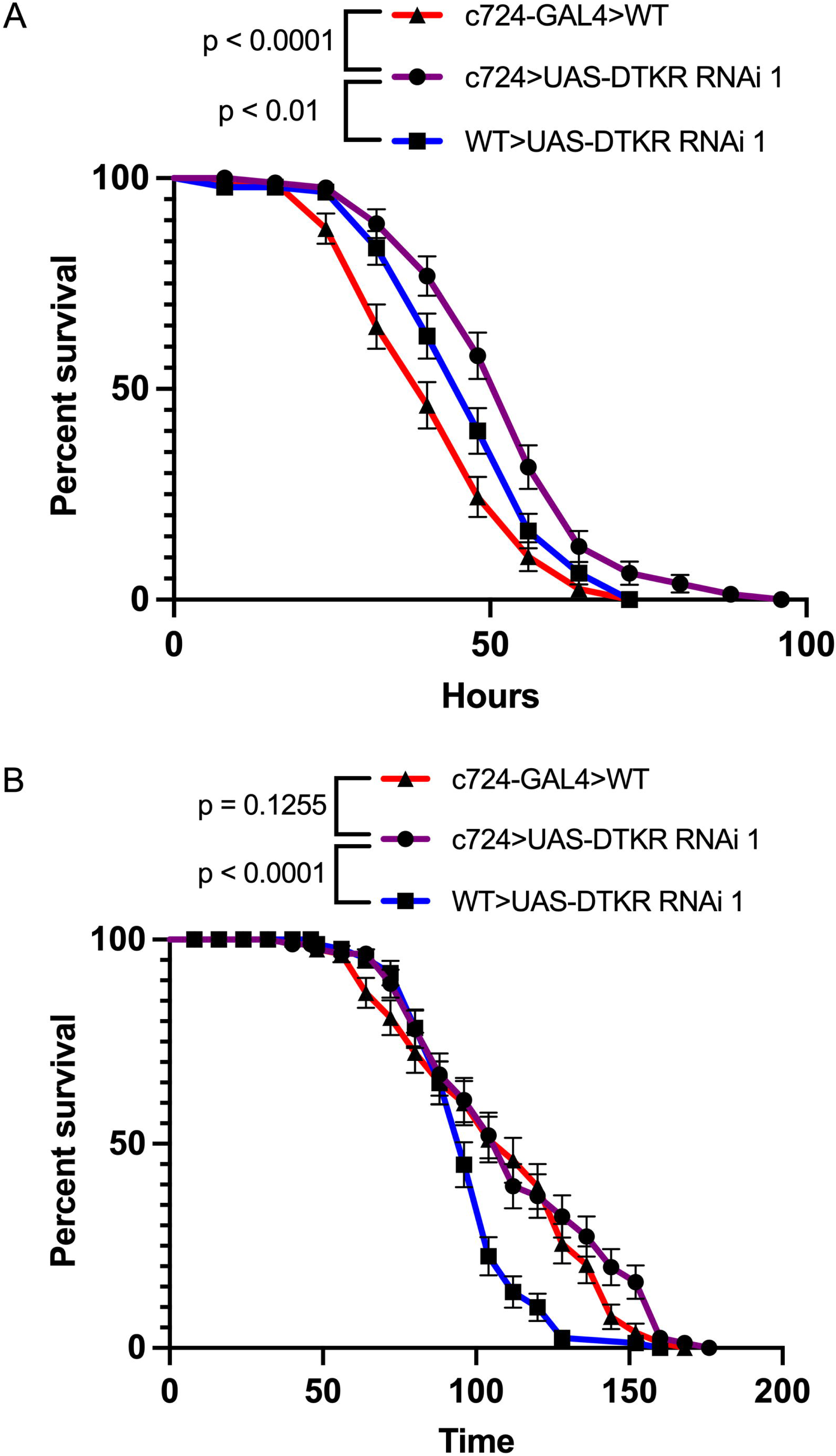
*DTKR* knockdown enhances desiccation stress survival while a role in starvation stress is unclear. (A) *DTKR*-RNAi flies driven by the stellate cell specific *c724*-GAL4 display significantly longer survival compared to the control flies under conditions of desiccation stress. (B) In response to starvation stress, *DTKR*-RNAi in tubule stellates cells increased the duration of survival compared to only one (of two) parental controls. (A) & (B) data are presented as survival curves assessed by Log-rank (Mantel-Cox) test after adjusting for multiple comparisons using the Bonferroni method. Significance levels (p value) for pairwise comparisons of survival curves are indicated in the legend above each figure.

## Discussion

In this study, we combined cell-type specific RNAi screening and various *ex vivo* and *in vivo* biological assays to provide evidence that DTKs are peptidergic hormones that interact with DTKR expressed in the MTs to contribute towards ionic and osmotic balance in the adult fruit fly, *D. melanogaster* (**Fig. 7**). Previous studies in other insects have indicated that tachykinins are not only myostimulatory on peripheral tissues but may also function as a diuretic factors increasing fluid secretion rates of isolated MTs (Schoofs et al., 1990a; Schoofs et al., 1990b; Siviter et al., 2000). However, since the MTs are not innervated, tachykinins elicit their activity on MTs by functioning as circulating hormones (Winther and Nässel, 2001). There have been attempts to localize the expression of the DTKR in the MTs, but these proved challenging in light of the contradictory evidence in the literature (Birse et al., 2006; Söderberg et al., 2011). Therefore, we attempted to functionally localize DTKR in specific cells of the MTs by utilizing the snRNA-seq analysis and GAL4/UAS bipartite system to knock down expression in the renal tubules using cell-specific drivers and characterize deficiencies in DTK-DTKR signaling by complementary *in vivo* and *ex vivo* bioassays.

**Fig. 7.**
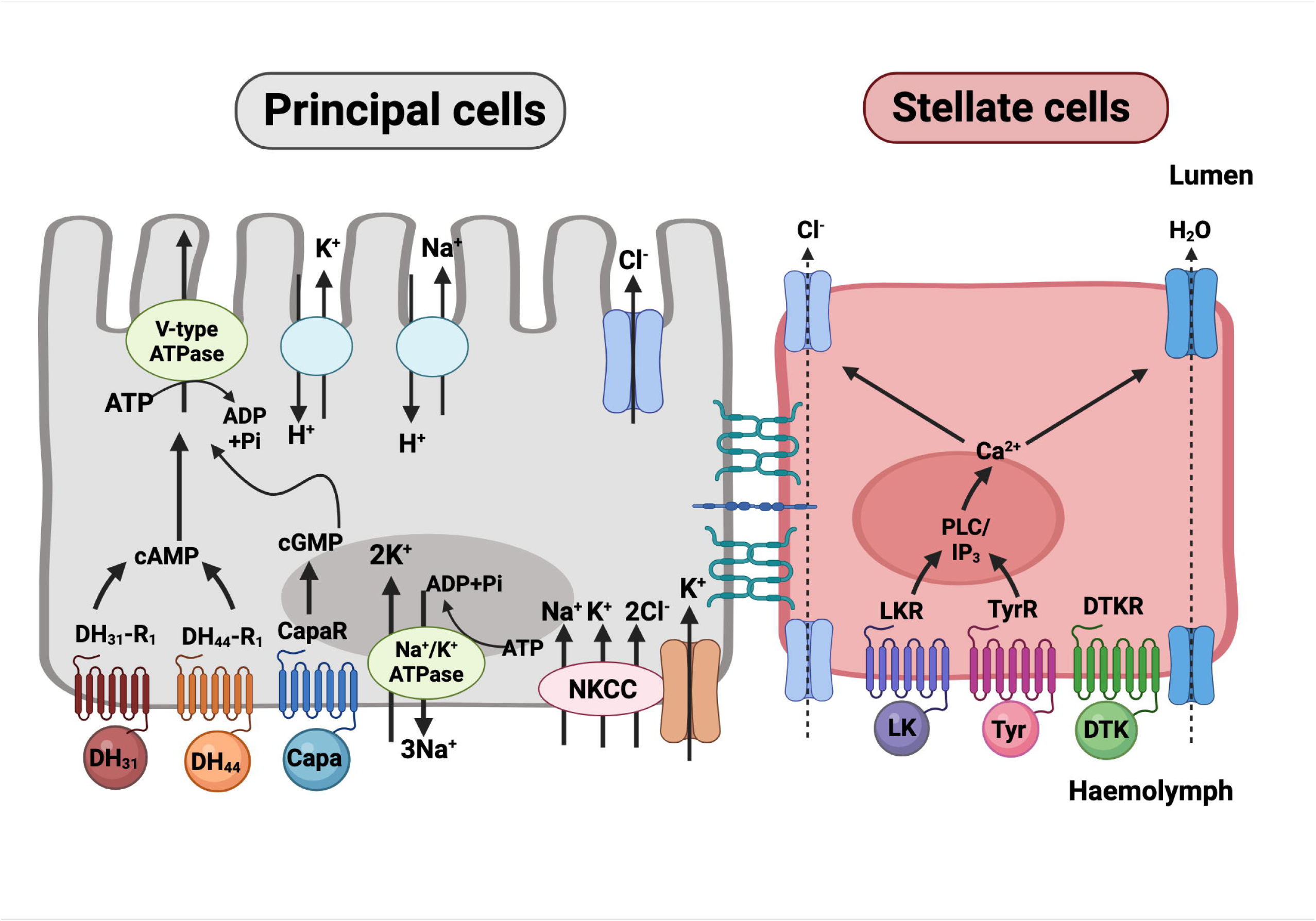
Schematic diagram depicting (neuro)endocrine factors targeting the principal and stellate cells in the *D. melanogaster* MTs. The principal cells are responsible for movement of the cations (Na^+^ and K^+^) driven by the V-type H^+^-ATPase localized on the apical membrane that creates a proton gradient. Neurohormones that target the principal cells includes: DH_31_, DH_44_, and Capa. The stellate cells are responsible for the movement of chloride ions driven by channels localized in the apical and basolateral membrane to mediate H_2_O transport via aquaporins. Diuretic hormones that stimulate transport in the stellate cells: LK, Tyr, and DTK.

Based on the snRNA-seq analysis, the expression of *TKR99D* was localized to the star-shaped stellate cells of the main segments of the MTs. In addition, we were able to confirm the localization along with other diuretic hormone receptors that are expressed in stellate cells. These include the receptors for leucokinin, DH_44_, and tyramine displaying high expression in the stellate cells of the MTs, which for the most part is consistent with previous studies (Cabrero et al., 2013; Cannell et al., 2016; Xu et al., 2022). Contrastingly, earlier evidence on the DH_44_ receptor suggested that it is expressed in principal cells where DH_44_ was reported to increase cAMP levels (Cabrero et al., 2002). Interestingly, however, principal cell driven RNAi of the DH_44_ receptor failed to abolish the diuretic activity of DH_44_ on isolated MTs (Cannell et al., 2016). The high expression of *TkR99D* in the stellate cells of the main tubule segment suggests that DTK/DTKR signaling may be an essential diuretic hormone for maintaining hydromineral balance by mediating transcellular transport of primarily chloride ions and water due to the high expression of chloride and water channels localized to the stellate cells (Cabrero et al., 2014; Cabrero et al., 2020; O’Donnell et al., 1998). Intriguingly, the fact that TkR99D-positive stellate cells display high expression of other diuretic hormone receptors, including the overlap in expression with the *Lkr*, suggest that these diuretic hormones may be functioning together to aid in transepithelial transport to maintain ion-water balance (Cabrero et al., 2020; Cannell et al., 2016).

Tachykinins in both vertebrates and invertebrates are potent stimulators inducing contractions of peripheral tissues, including the hindgut, oviduct, and the MTs in insects (Coast, 1998; Schoofs et al., 1990a; Schoofs et al., 1990b; Siviter et al., 2000; Winther et al., 1998). Since previous studies demonstrated that tachykinins in other insect species act as diuretic hormones (Johard et al., 2003; Skaer et al., 2002), we aimed to determine whether this peptidergic signaling system plays a similar role in the fruit fly. Indeed, we observed that DTKs are also potent stimulators of fluid secretion by isolated MTs whereby a dose-dependent increase in fluid secretion rate is observed, with an EC_50_ in the nanomolar range. This physiological dose of DTK is consistent with previous studies that have investigated other diuretic factors like tyramine and leucokinin that target stellate cells in the MTs inducing a rapid increase in fluid transport (Cabrero et al., 2013; Terhzaz et al., 1999). Relatedly, previous studies have highlighted the diuretic role of different isoforms of insect tachykinin-related peptides at various concentrations, including in the pharate adult *Manduca sexta*, where concentrations in the range of 0.01-1 µM elicit diuretic activity on isolated MTs (Skaer et al., 2002).

The next question we addressed in this study was, where are DTKs specifically acting in the MTs to elicit their diuretic activity? Our findings indicate that DTKR is expressed in the main segment stellate cells and not in principal cells of the MTs. This conclusion is based on multiple lines of evidence including *DTKR* knock down driven by the stellate cell driver which had significantly lower *DTKR* transcript abundance whereas no change in transcript level was observed when knock down was attempted using the principal cell driver. These findings are consistent with our snRNA-seq analysis using a recent dataset that characterized the cell types of the adult fly MTs (Xu et al., 2022). Specifically, the star-shaped main segment stellate cells were found to express *DTKR* in support of the notion that these cells in the MTs are the target of circulating DTKs. The results also corroborate the initial study that immunolocalized DTKR in the bar-shaped stellate cells in the larval fruit fly MTs (Birse et al., 2006). Not only was DTKR transcript abundance affected, but stellate cell driven knock down also significantly reduced fluid secretion rates stimulated by DTK-1 by more than 80%. Comparatively, MTs with principal cell driven knock down of DTKR had normal DTK-1 stimulated secretion rates. These results align with data from earlier studies that found knock down of receptors for diuretic factors in the stellate cells resulted in reduced stimulated secretion rates compared to control fly lines (Cabrero et al., 2013; Cannell et al., 2016; Feingold et al., 2019).

Altogether the data implies that DTK is a diuretic hormone that targets DTKR in the tubule stellate cells, which like leucokinin and tyramine (Cabrero et al., 2013; Cannell et al., 2016; Feingold et al., 2019; Terhzaz et al., 1999), contribute towards the regulation of hydromineral balance in the insect. In light of the diuretic activity of DTK on the stellate cells of MTs and considering DTKR downstream signaling involves increases in Ca^2+^ (Birse et al., 2006; Im et al., 2015), it may trigger a second messenger pathway similar to the other diuretics targeting the stellate cells to stimulate fluid secretion. The importance of calcium and confirmation of its involvement in eliciting DTK-DTKR signaling in the tubule stellate cells awaits further investigation, but such signaling has been deemed important for modulating nociceptive sensitization in *Drosophila* (Im et al., 2015).

Besides establishing its role as a diuretic hormone leading to increased fluid secretion, we were particularly interested in discerning how this diuretic activity is achieved given water cannot be moved across the tubule epithelia without a favourable osmotic gradient, provided largely through passage of inorganic ions. To further characterize the mechanism by which DTK elicits diuretic action on the MTs, we measured Na^+^, K^+^, and Cl^−^ transport by isolated MTs. The *ex vivo* bioassay results revealed that DTK stimulates Na^+^, K^+^ & Cl^−^ transport, which in turn promotes osmotically-obliged water leading to fluid secretion from the tubules. However, the transport rate data alone does not confirm whether the primary effect of DTK involves stimulating Cl^−^ transport. Through additional studies involving electrophysiological recordings, it may be established that DTK stimulation of ion transport is similar to actions of kinin-related neuropeptides in the mosquito *Ae. aegypti*, which increases Cl^−^ conductance driving transepithelial transport of Na^+^ and K^+^ (Hayes et al., 1994; Pannabecker et al., 1993). Similarly, the endocrine control of chloride conductance by MTs was shown in *Drosophila*, where leucokinin was found to stimulate chloride flux transcellularly through the chloride channel encoded by CG31116 (Cabrero et al., 2014). Similarly, the biogenic amine tyramine, which is produced endogenously by MTs and thus elicits its control in a paracrine fashion, acts identically to leucokinin leading to increased chloride permeability via stellate cells (Blumenthal, 2003; Cabrero et al., 2013). Therefore, DTK-1 stimulated fluid secretion may likely involve the increased chloride flux, which ultimately leads to depolarization (or collapsing) of the transepithelial membrane potential across the tubules (O’Donnell et al., 1998). If this is the case, then the transport of chloride in the stellate cells is coupled with the active transport of cations in the principal cells, which in turn increases transepithelial transport across the tubules upon DTK stimulation with the help of transporters localized in the principal cells, including the Na^+^-K+-2Cl^−^ (NKCC) and Na^+^-K^+^-ATPase (NKA) (Blumenthal, 2003; Cabrero et al., 2013; Linton and O’Donnell, 1999; Pannabecker et al., 1993; Rodan et al., 2012). In addition, the transport of cations and anions may drive the movement of water via aquaporins localized in the stellate cells or the entomoglyceroporins in principal cells (Cabrero et al., 2020). Future investigations should look at the activity and involvement of specific channels and transporters localized in the stellate cells when MTs are stimulated by DTK peptides. For example, previous studies have shown that stellate cell targeting diuretic factors (leucokinin and tyramine) utilize different chloride channels (*ClC-a* and *secCl*) either on the apical or basolateral membranes of the stellate cells in the MTs of the fruit fly (Cabrero et al., 2014; Feingold et al., 2019). The current results involving DTK-1 stimulated diuretic activity appears to demonstrate changes in transepithelial transport. In particular, both chloride concentration and transport rate increased significantly in response to DTK-1 in wildtype flies, corroborating that this peptidergic hormone is directly targeting the stellate cells having abundant chloride channels necessary for stimulated fluid secretion, and osmotically drawn water transport in the same (stellate) and adjacent (principal) cells (Cabrero et al., 2020). Moreover, knockdown of DTKR specifically in the tubule stellate cells mainly affects the transport rate and not the concentration of Na^+^ and K^+^ while affecting both the transport rate and concentration of Cl^−^ ions. Therefore, the dramatic reduction in DTK-1 stimulated fluid secretion rates in *DTKR* knockdown flies is likely due to a drop in transepithelial ion transport resulting in reduced osmotic pressure that lowers the driving force for water transport. Additionally, changes in the chloride concentration in the absence of changes in cation concentrations could impact pH of the secreted fluid and, relatedly, have implications for whole animal pH balance given Cl^−^ transport in dipteran MTs is linked to the other major anion, bicarbonate, via an exchanger localized to stellate cells (Piermarini et al., 2010).

Considering the functional consequences of DTKR knockdown on isolated MTs *ex vivo*, we questioned whether this reduced DTK-DTKR signaling involving the Malpighian ‘renal’ tubules alone might impact adult fly survival during starvation or desiccation stress, raising the notion that such signaling plays an important role in whole organismal physiology. Indeed, our data indicates that *DTKR* knockdown in the MTs prolonged survival of flies during desiccation stress, while with respect to starvation stress, we were unable to draw the same conclusion since knockdown flies survived longer when compared to only one of two parental controls. Despite converging on the same renal cell type and driving chloride transport, one important difference between leucokinin and DTK signaling is that knockdown of LKR in stellate cells improves starvation tolerance but not desiccation tolerance (Cannell et al., 2016), which represents the opposite phenotype following *DTKR* knockdown. DTKs are extensively distributed and synthesized in the central nervous system including over 150 neurons in the adult fly brain, although it has been argued that this neural origin likely does not result in hormonal release since most DTK producing cells in the brain are interneurons (Nässel et al., 2019). A subset of ten DTK producing neurons in adult *D. melanogaster* brain (ITPn cells) are neurosecretory cells, that co-express ITP and sNPF (Kahsai et al., 2010). Since ITP from ITPn functions as an anti-diuretic hormone, it seems counter-intuitive for diuretic DTK to be co-released into the circulation (Gera et al., 2024). In support of this hypothesis, it has been argued that it is unlikely that DTK (or sNPF) derived from ITPn cells are released as circulating hormones but instead might act locally as neuromodulators in other brain regions or presynaptically on ITPn axon terminals (Kahsai et al., 2010; Nässel et al., 2019). Interestingly, most recent evidence suggests that this speculation of local release might be incorrect since ITP producing neurosecretory cells do not form synapses in the brain (Gera et al., 2024). On the other hand, strong evidence supports that the midgut is responsible for endocrine action of DTKs in circulation as intestinal release has been demonstrated in other insects, including locusts and cockroaches (Winther and Nässel, 2001). Circulating DTKs acting as hormones target, as confirmed herein, DTKR expressed in stellate cells of the MTs triggering ion and fluid transport, which may lead to greater water loss from the organism. Relatedly, *DTKR* knockdown solely in the stellate cells of the MTs improved the survival during desiccation stress suggesting that these animals are more tolerant to dehydration. These results are consistent with an earlier study where knockdown of diuretic hormones in the fruit fly, including the leucokinin hormone, resulted in increased survival during desiccation stress (Zandawala et al., 2018). Interestingly, internal stimuli may be integrated differently in relation to DTK signaling and it may depend on the source of this neuropeptide since desiccation stress led to increased water loss when *Dtk* was diminished in a subset of DTK producing neurons in the brain, resulting in increased desiccation sensitivity (Kahsai et al., 2010), the opposite phenotype to what we observed when DTKR was reduced exclusively in stellate cells. This could be explained by considering brain-derived DTKs in ITPn (Gera et al., 2024; Kahsai et al., 2010) might affect other hormones acting on the MTs and other organs controlling hydromineral homeostasis, whereas DTKR knockdown exclusively in stellate cells reveals a more direct phenotype of altered DTK/DTKR signaling.

In summary, our data confirm that DTKs regulate fluid and ion transport by acting on the stellate cells of the MTs in the adult fruit fly, *D. melanogaster*. Although the source of circulating DTKs acting on the MTs remains unconfirmed in flies, they are likely released from enteroendocrine cells in the midgut as established in other insects, including cockroach and locust (Siviter et al., 2000; Winther and Nässel, 2001). DTKs are localized in enteroendocrine cells in the midgut and may regulate lipid metabolism, especially during times of starvation (Song et al., 2014). Indeed, this earlier study established that tachykinins derived from the midgut regulate feeding-related physiological processes, while tachykinins derived from the nervous system govern behaviour (Song et al., 2014). Overall, this suggests that tachykinins could be released in response to physiological stress in the fly and future studies should confirm the source of hormonal DTKs along with the specific stimuli that control their release. To conclude, DTKs are essential for osmoregulation in *D. melanogaster* given our findings reveal these peptide hormones contribute towards organismal ion-water balance by targeting DTKR expressed in stellate cells of the MTs to trigger diuresis. DTK/DTKR diuretic signaling may contribute towards hydromineral homeostasis and nutritional balance, considering its earlier documented involvement in increasing resistance to starvation stress (Söderberg et al., 2011), these two pathways may converge to coordinate metabolic and osmoregulatory demands of the fly, particularly in response to nutritional and desiccation stress.

## Acknowledgements

The authors would like to thank Dr. Kenneth Halberg (University of Copenhagen) and Dr. Laura Nilson (McGill University) for providing the following fly lines: *uro*-GAL4 and *c724*-GAL4, respectively. The authors are grateful to Dr. Andrew Donini (York University) for providing critical insight and suggestions with respect to analysis and interpretation of electrophysiological data. This work was supported by a Natural Sciences and Engineering Research Council of Canada (NSERC) Discovery Grant and an Ontario Ministry of Research Innovation Early Researcher Award to JPP.

